# ApoplastP: prediction of effectors and plant proteins in the apoplast using machine learning

**DOI:** 10.1101/182428

**Authors:** Jana Sperschneider, Peter N. Dodds, Karam B. Singh, Jennifer M. Taylor

## Abstract

The plant apoplast is integral to intercellular signalling, transport and plant-pathogen interactions. Plant pathogens deliver effectors both into the apoplast and inside host cells, but no computational method currently exists to discriminate between these localizations. We present ApoplastP, the first method for predicting if an effector or plant protein localizes to the apoplast. ApoplastP uncovers features for apoplastic localization common to both effectors and plant proteins, namely an enrichment in small amino acids and cysteines as well as depletion in glutamic acid. ApoplastP predicts apoplastic localization in effectors with sensitivity of 75% and false positive rate of 5%, improving accuracy of cysteine-rich classifiers by over 13%. ApoplastP does not depend on the presence of a signal peptide and correctly predicts the localization of unconventionally secreted plant and effector proteins. The secretomes of fungal saprophytes, necrotrophic pathogens and extracellular pathogens are enriched for predicted apoplastic proteins. Rust pathogen secretomes have the lowest percentage of apoplastic proteins, but these are highly enriched for predicted effectors. ApoplastP pioneers apoplastic localization prediction using machine learning. It will facilitate functional studies and will be valuable for predicting if an effector localizes to the apoplast or if it enters plant cells. ApoplastP is available at http://apoplastp.csiro.au.

## Introduction

Pathogenic microbes such as bacteria, fungi, oomycetes, and nematodes colonize and infect plant cells and cause devastating diseases and crop losses. The extracellular matrix of plant tissues is known as the apoplast and is integral to plant physiology, signalling and defence against plant-pathogenic microbes. Initial contact between plant and pathogens is made in the apoplast and early interactions determine if a pathogen is able to colonize its host. Plant cell surface localized pattern recognition receptors (PRRs) recognize conserved pathogen molecules known as pathogen-associated molecular patterns (PAMPs) or microbe-associated molecular patterns (MAMPs) and launch initial defence responses (Dodds & Rathjen, 2010). Activation of PRR signalling leads to PAMP-triggered immunity (PTI) with rapid accumulation of antimicrobial compounds and proteins such as proteinases, chitinases, glucanases and enzyme inhibitors that damage pathogen structures and molecules (Lo Presti *et al.*, 2015). In turn, plant pathogens secrete effectors that alter host-cell structure and function, thereby facilitating infection and/or triggering defence responses (Kamoun, 2006). Apoplastic effectors can function as enzyme inhibitors, scavenge molecules that trigger plant immune responses and protect pathogen infection structures such as hyphae from recognition (Lo Presti *et al.*, 2015). Some pathogens also deliver cytoplasmic effectors into plant cells to target intracellular processes. Both apoplastic and cytoplasmic effectors can also be recognized by either membrane bound or intracellular plant receptors to trigger defence responses often known as effector-triggered immunity (ETI) (Stotz *et al.*, 2014).

Plant pathogens have evolved various strategies to deliver cytoplasmic effectors intracellularly. Biotrophic and hemibiotrophic pathogens must suppress host defences as they feed on living plant cells, whereas necrotrophic pathogens feed and grow on dead or dying plant tissue and trigger host cell death as a colonization strategy. Some pathogens can directly penetrate plant tissue through specialized infection structures and deliver cytoplasmic effectors into the plant cell. For example, the hemibiotrophic fungal pathogens *Magnaporthe oryzae* and *Colletotrichum higginsianum* enter plant cells through melanized appressoria, whereas the biotrophic fungal pathogen *Ustilago maydis* uses non-melanized appressoria to invade host cells (Giraldo & Valent, 2013). Rust fungi, powdery mildews and oomycetes can form dedicated feeding structures called haustoria that act as sites of effector delivery to the plant cell cytoplasm (Garnica *et al.*, 2014). Other fungal pathogens such as *Cladosporium fulvum, Zymoseptoria tritici, Leptosphaeria maculans* and *Venturia inaequalis* colonize plants extracellularly and rely on apoplastic effectors to target basal apoplastic host defence components (Stotz *et al.*, 2014; Zhong *et al.*, 2017).

The diversity of plant pathogen effectors poses a challenge for their prediction from genomic sequences. Bacterial cytoplasmic effectors are generally predicted using machine learning methods based on conserved host delivery mechanisms such as the type III secretion system (McDermott *et al.*, 2011). Cytoplasmic oomycete effectors are commonly predicted based on the presence of conserved N-terminal sequence motifs, but this analysis is typically restricted to RxLR or Crinkler effector families (Bhattacharjee *et al.*, 2006). Effector prediction in fungal pathogens is complicated by the lack of conserved sequence features or motifs. User-driven selection of proteins with a small size and a high number of cysteines is commonly used to mine fungal secretomes for effectors, but suffers from poor accuracy especially for cytoplasmic effectors (Sperschneider *et al.*, 2015a). Fungal effector prediction can benefit from including evidence of diversifying selection (Guyon *et al.*, 2014; Sperschneider *et al.*, 2014) or the genomic context of the gene for pathogens that preferentially harbour effectors in genomic regions with higher evolutionary rates (Raffaele & Kamoun, 2012). Whilst such methods are powerful, they only capture a subset of the effector repertoire as these are not necessarily universal signals for both apoplastic and cytoplasmic effectors. By contrast, a data-driven machine learning classifier can learn ‘effector rules’ from positive and negative training examples without having to apply user-chosen thresholds, and this was exemplified in the first machine learning classifier for fungal effector prediction called EffectorP (Sperschneider *et al.*, 2016). However, EffectorP is not able to distinguish between apoplastic and cytoplasmic fungal effectors.

Machine learning methods can classify proteins by recognising patterns when informative sequence homologies or motifs are missing, and are thus promising for predicting effectors and their localisation. A recent method called LOCALIZER has improved prediction ability for targeting signals to plant chloroplasts, mitochondria and nuclei in effectors (Sperschneider *et al.*, 2017). Signal peptide prediction tools such as SignalP (Petersen *et al.*, 2011) and Phobius (Kall *et al.*, 2007) as well as plant subcellular localization predictors such as WoLF PSORT (Horton *et al.*, 2007) or YLoc (Briesemeister *et al.*, 2010) can predict extracellular localization, but not apoplastic localization specifically. For extracellular pathogens, accurate prediction of apoplastic effector candidates is important for prioritizing host-recognized Avr effectors for experimental validation. For intracellular pathogens, effector candidates with a predicted signal peptide but with non-apoplastic localization are prime candidates for prioritizing Avr effectors for experimental validation. In oomycetes, the presence of the RxLR motif or the Crinkler domain have been used as proxies for predicting host-translocation and thus their intracellular localization (Petre & Kamoun, 2014). However, recent evidence suggests that the RxLR motif might play a role in intracellular processing before secretion (Wawra *et al.*, 2017). No conserved sequence motif with a role in host translocation has thus far been found for fungal pathogens (Sperschneider *et al.*, 2015a) that can be utilized to predict cytoplasmic localization for effectors. Taken together, for both plant and pathogen proteins, no computational method currently exists to determine apoplastic localization despite its importance in plant-pathogen interactions and its value in guiding experimental validation.

Apoplastic proteins can be identified through microscopic analyses or apoplastic proteomics, however both are technically challenging (Doehlemann & Hemetsberger, 2013; Delaunois *et al.*, 2014). The first challenge for *in planta* proteomics is the collection of sufficient apoplastic material without causing cell wall damage and thus contamination with cytoplasmic proteins. Alternatively, *in vitro* experiments can limit contamination with cytoplasmic proteins, but only have partial ability to characterize apoplastic proteins involved in plant-pathogen interactions (Jung *et al.*, 2008). There is increasing evidence that apoplastic proteins can be secreted unconventionally (Delaunois *et al.*, 2014) and these cannot be detected in the apoplastic proteome by signal peptide prediction tools such as SignalP (Emanuelsson *et al.*, 2007). Currently, the only prediction tool for unconventionally secreted proteins is SecretomeP (Bendtsen *et al.*, 2004a), but it has been trained on mammalian sequences and is not recommended for use on plants or pathogens (Lonsdale *et al.*, 2016). Taken together, the technical challenges of proteomics and microscopic analyses as well as the lack of bioinformatics tools for apoplastic localization prediction has limited progress in our understanding of early plant-pathogen interactions in the apoplast, in the identification of alternative secretion pathways and in the ability to discriminate between apoplastic and cytoplasmic effectors.

## Description

### Training and evaluation of the machine learning classifiers

Literature searches were performed to collect apoplastic and cytoplasmic effectors with experimental support for both the training and independent test sets (Table 1, FASTA sequences available at http://apoplastp.csiro.au/data).

**Table 1:** Effectors used in the training and independent test sets.

| Data set | Used in training | Used in independent testing |
| --- | --- | --- |
| **Apoplastic effectors with experimental validation** |  |  |
| Fungal | AvrLm4-7, Ave1, Avr9, Avr4, Avr4E, Avr2, Avr5, Ecp2, Ecp1, Ecp5, Ecp4, Bas4, MC69, Slp1, NIP1, Pep1, Pit2, Mg3Lysm | Ecp6, AvrLm6, AvrLm1, AvrLm11, NEP1, ToxB, Msp1, AvrStb6, Cgfl, PstSCR1, CfTom1 |
| Oomycete | GIP1, EPI1, PcF, EpiC1, CBEL, NPP1 | INF1, GIP2, EPI10, INF2A, INF2B, EPIC2A, EPIC2B, PsXEG1, CBEL, Ecpic1, OPEL, GP42, NIP1 |
| Nematode | - | Asp2, Crt-1, Map-1, VAP1, HYP-3, Sp12, CLE1, CLE2 |
| **Cytoplasmic effectors with experimental validation** |  |  |
| Fungal | - | AvrM, AvrL567, AvrP123, AvrP4, RTP1, PGTAUSPE-10-1, Avra10, Six3, Six6, Avr-Pita1, Pwl1, Avr-Pia, Bas1, AvrPiz-t, Avr1-CO39, Avr-Pii, Avr-Pik, Bas107, See1, Cmu1, Tin2, ToxA, SP7, MISSP7, Bas162, AvrM14, AvrL2-A, CgEP1, VdSCP7 |
| Oomycete | - | Atr13, Avr3a, Avr1B-1, Atr1, Avh5, Avh241, Avr1l, Avr1d, Avrblb1, Avrblb2, Avr2, Crn1, Crn2, Crn8, Crn15, Crn16, Crn63, Crn115, Atr5, PexRD2, Atr1, Avr3b, PITG_03192, Avr1, Pslsc1, Atr39-1, Avh18a1, Avr1a, Avr3a, HaAtr1, PiAvr2, PiAvr3b, PiAvr4, PiAvrVnt1, PsAvr3b, PsAvr3c, PsAvr4/6, SNE1 |

As a positive training set, 349 apoplastic plant proteins (retrieved from UniProt: taxonomy: *Viridiplantae* locations:(location:apoplast) AND reviewed:yes) as well as 24 apoplastic, experimentally validated effectors from fungal and oomycete pathogens from the literature (Table 1) were collected. Only sequences > 50 aas and starting with ‘M’ were considered. The 373 sequences were homology-reduced as follows. First, a sequence was randomly picked and added to the homology-reduced set if it did not share significant sequence similarity with another sequence already present in the homology-reduced set. Significant sequence similarity was assessed using phmmer (Finn *et al.*, 2011) with a bit score threshold of larger than 100. This resulted in a positive training set of 84 proteins (FASTA sequences available at <u>http://apoplastp.csiro.au/data</u>).

Non-extracellular plant proteins from the UniProt database (chloroplast, cytoplasm, membranes, mitochondria, nucleus) were used as the negative training set (taxonomy: “Viridiplantae [33090]” and supported by experimental evidence: chloroplast (“Plastid [SL-0209]”, “Chloroplast [SL-0209]”); cytoplasm (“Cytoplasm [SL-0086]”); membranes (“membrane”); mitochondria (“Mitochondrion [SL-0173]”); nucleus (“Nucleus [SL-0191]”)). These 1,950 sequences were homology-reduced and this resulted in a negative training set of 1,773 proteins. For each protein, the feature vector used the following features calculated with pepstats (Rice *et al.*, 2000): percentages of amino acids (A, C, D, E, F, G, H, I, K, L, M, N, P, Q, R, S, T, V, W, Y) and percentages of amino acid classes (tiny, small, aliphatic, aromatic, nonpolar, polar, charged, basic, acidic) in the sequence, total number of cysteines in the sequence, protein net charge, isoelectric point, grand average of hydropathicity (GRAVY) as well as the protein instability index and protein aromaticity calculated using ProtParam (Gasteiger *et al.*, 2005). As ProtParam does not allow ambiguous amino acids as input, we replaced these with randomly selected respective amino acids (B replaced with D or N; Z replaced with E or Q, X replaced with any amino acid).

Weka 3.8.1 was used to train machine learning classifiers (Frank, 2016). For the Random Forest classifier, proteins with probability > 0.55 were classified as apoplastic. Weka’s CorrelationAttributeEval + Ranker method was used to find the most discriminative features for classification.

In the evaluation, a true positive (TP) is an apoplastic protein that is correctly predicted as an apoplastic protein and a false positive (FP) is a non-apoplastic protein incorrectly predicted as an apoplastic protein. A true negative (TN) is a non-apoplastic protein that is correctly predicted as a non-apoplastic protein and a false negative (FN) is an apoplastic protein incorrectly predicted as a non-apoplastic protein. Sensitivity 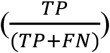 is defined as the proportion of positives that are correctly identified whereas specificity 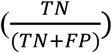 is the proportion of negatives that are correctly identified. Precision (positive predictive value, PPV) is a measure which captures the proportion of positive predictions that are true 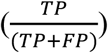 Both accuracy 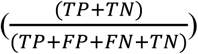and the Matthews Correlation Coefficient 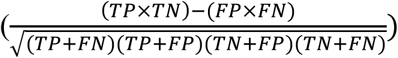 were also used to evaluate the overall performance of the method. The MCC ranges from −1 to 1, with scores of −1 corresponding to predictions in total disagreement with the observations, 0 to random predictions and 1 to predictions in perfect agreement with the observations. The receiver operating characteristic (ROC) curve is drawn by plotting sensitivity against (1 – specificity) and the area under the curve (AUC) can be interpreted as the probability that a classifier will rank a randomly chosen apoplastic protein higher than a randomly chosen non-apoplastic protein. Therefore, a perfect classifier achieves an AUC of 1.0, whereas a random classifier achieves an AUC of only 0.5.

For the evaluation, we collected plant and fungal proteins that have been experimentally shown to localize to the ER, Golgi, vacuole or contain transmembrane domains (taxonomy:"Fungi [4751]" locations:(location:"Endoplasmic reticulum [SL-0095]" evidence:experimental) AND reviewed:yes; locations:(location:golgi evidence:experimental) AND reviewed:yes; taxonomy: locations:(location:"Vacuole [SL-0272]" evidence:experimental) AND reviewed:yes; transmembrane AND reviewed:yes; taxonomy:"Viridiplantae [33090]" locations: (location:"Endoplasmic reticulum [SL-0095]" evidence:experimental) AND reviewed:yes; locations:(location:golgi evidence:experimental) AND reviewed:yes; locations: (location:"Vacuole [SL-0272]" evidence:experimental) AND reviewed:yes; transmembrane AND reviewed:yes) and do not have the terms ‘extracellular’, ‘secreted’, ‘cytoplasm’ or ‘nucleus’ as additional subcellular localization or ‘extracellular’ in the description of the UniProt entry. We also collected extracellular mammalian proteins from UniProt (taxonomy:"Mammalia [40674]" locations:(location:extracellular evidence:experimental)). SignalP 4.1 (Petersen *et al.*, 2011) was run on all these sets and only proteins that have a predicted signal peptide were kept.

All plots were produced using ggplot2 (Wickham, 2009) and statistical significance was assessed with *t*-tests using the ggsignif package (https://cran.r-project.org/web/packages/ggsignif/index.html). Significance thresholds according to t-test are NS: not significant, * < 0.05, ** < 0.01 and *** < 0.001.

### Secretome predictions, effector predictions and sequence motif searches

The following fungal and oomycete genomes were collected: *Hyaloperonospora arabidopsidis* (Baxter *et al.*, 2010); *Albugo laibachii* (Kemen *et al.*, 2011); *Melampsora laricis-populina* and *Puccinia graminis* f. sp. *tritici* (Duplessis *et al.*, 2011); *Melampsora lini* (Nemri *et al.*, 2014); *Puccinia triticina* (Puccinia Group Sequencing Project); *Puccinia striiformis* f. sp. *tritici* PST-130 (Cantu *et al.*, 2011); *Blumeria graminis* f. sp. *hordei* (Spanu *et al.*, 2010); *Blumeria graminis* f. sp. *tritici* (Wicker *et al.*, 2013); *Ustilago maydis* (Kämper *et al.*, 2006); *Venturia pirina* (Cooke *et al.*, 2014); *Venturia inaequalis* (Deng *et al.*, 2017); *Cladosporium fulvum* (de Wit *et al.*, 2012); *Phytophthora infestans* (Haas *et al.*, 2009); *Phytophthora capsici* (Lamour *et al.*, 2012); *Phytophthora sojae* and *Phytophthora ramorum* (Tyler *et al.*, 2006); *Fusarium graminearum* (Cuomo *et al.*, 2007), *Fusarium oxysporum* f. sp. *lycopersici* and *Fusarium oxysporum* 47 (Ma *et al.*, 2010); *Leptosphaeria maculans* (Rouxel *et al.*, 2011); *Magnaporthe oryzae* (Dean *et al.*, 2005); *Zymoseptoria tritici* (Goodwin *et al.*, 2011); *Verticillium dahliae* (Klosterman *et al.*, 2011); *Colletotrichum higginsianum* (O’Connell *et al.*, 2012); *Pythium ultimum* (Levesque *et al.*, 2010); *Stagonospora nodorum* (Hane *et al.*, 2007); *Botrytis cinerea* and *Sclerotinia sclerotiorum* (Amselem *et al.*, 2011); *Rhizoctonia solani* AG8 (Hane *et al.*, 2014); *Pyrenophora tritici-repentis* (Manning *et al.*, 2013); *Penicillium digitatum* (Marcet-Houben *et al.*, 2012); *Laccaria bicolor* (Martin *et al.*, 2008); *Amanita muscaria*, *Hebeloma cylindrosporum*, *Laccaria amethystina*, *Paxillus involutus*, *Paxillus rubicundulus*, *Piloderma croceum*, *Pisolithus microcarpus*, *Pisolithus tinctorius*, *Scleroderma citrinum*, *Sebacina vermifera*, *Suillus luteus* and *Tulasnella calospora* (Kohler *et al.*, 2015); *Aspergillus flavus* (Arnaud *et al.*, 2012); *Tremella mesenterica*, *Coniophora puteana*, *Dacryopinax sp*., *Fomitopsis pinicola, Gloeophyllum trabeum, Wolfiporia cocos*, *Dichomitus squalens, Fomitiporia mediterranea, Punctularia strigosozonata, Stereum hirsutum* and *Trametes versicolor* (Floudas *et al.*, 2012); *Rhodotorula graminis* (Firrincieli *et al.*, 2015); *Batrachochytrium dendrobatidis* (https://www.broadinstitute.org/fungal-genome-initiative/batrachochytrium-genome-project); *Ashbya gossypii* (Gattiker *et al.*, 2007); *Taphrina deformans* (Cisse *et al.*, 2013); *Pichia stipitis* (Jeffries *et al.*, 2007); Sacch*aromyces cerevisiae* (Goffeau *et al.*, 1996); *Hysterium pulicare* (Ohm *et al.*, 2012); *Coprinus cinereus* (Stajich *et al.*, 2010); *Aspergillus oryzae* (Machida *et al.*, 2005); *Aspergillus niger* (Andersen *et al.*, 2011); *Neurospora crassa* (Galagan *et al.*, 2003); *Agaricus bisporus var bisporus* (Morin *et al.*, 2012); *Rhodosporidium toruloides* (Zhu *et al.*, 2012). Secretome predictions of fungal and oomycete genomes was done using SignalP 3 (Bendtsen *et al.*, 2004b), TMHMM (Krogh *et al.*, 2001) and TargetP (Emanuelsson *et al.*, 2000) as described in Sperschneider *et al.* (2015b). Effector candidates were predicted using EffectorP 1.0 (Sperschneider *et al.*, 2016). MEME motif searches (Bailey *et al.*, 2009) were run on the EffectorP 1.0 predicted apoplastic and non-apoplastic effector candidates after sequence homology reduction. MEME was run with the parameters –protein –nmotifs 5 –mod oops.

### *Puccinia graminis* f. sp. *tritici* (*Pgt*) 21-0 differential expression analysis

Reads for germinated spores and haustorial tissue (100bp paired-end) were obtained from NCBI BioProject PRJNA253722 (Upadhyaya *et al.*, 2014). These were adapter-trimmed using trimgalore with default parameters (https://www.bioinformatics.babraham.ac.uk/projects/trim_galore/). Reads were aligned to the *Pgt* 21-0 genomes using STAR with default parameters (Dobin *et al.*, 2013). FeatureCounts (Liao *et al.*, 2014) was used to obtain a read count matrix. The DESeq2 package was used for differential expression analysis of the *Pgt* 21-0 gene set (Love *et al.*, 2014). Genes showing differential expression (adjusted *p-*value padj < 0.1) in haustorial tissue versus germinated spores were selected at logFC thresholds of -1.0, 1.0, -10 and 10.

## Results

### An enrichment in cysteines is a feature of apoplastic fungal and oomycete effectors, but not of apoplastic plant proteins

The plant apoplast is a harsh physiological environment rich in degradative proteases (Kamoun, 2006; Lo Presti *et al.*, 2015) and is likely to impose particular stability constraints on apoplastic proteins. We first investigated if a small size and high cysteine content, as routinely used as criteria for fungal effector prediction (Stergiopoulos & de Wit, 2009; Sperschneider *et al.*, 2015a), is sufficient for predicting apoplastic localization. First, we compared 29 experimentally validated apoplastic fungal effectors to 29 experimentally validated cytoplasmic fungal effectors (Table 1). We observed no significant differences between the two groups in terms of sequence length, but we found a significantly higher percentage of cysteines as well as a higher total number of cysteines for apoplastic fungal effectors (Fig. 1A). We then tested simple classifiers using different thresholds for cysteine content and found that this resulted in high false positive rates of 19.2% to 43.7%. This suggests that thresholds for cysteine content do not allow for highly accurate discrimination of apoplastic effectors from cytoplasmic effectors in fungi (Table 2). For example, a small size and high cysteine content are also found in intracellular fungal effectors such as the *Melampsora lini* effectors AvrP123 (117 aas, 11 cysteines) and AvrP4 (95 aas, 7 cysteines).

**Fig. 1:**
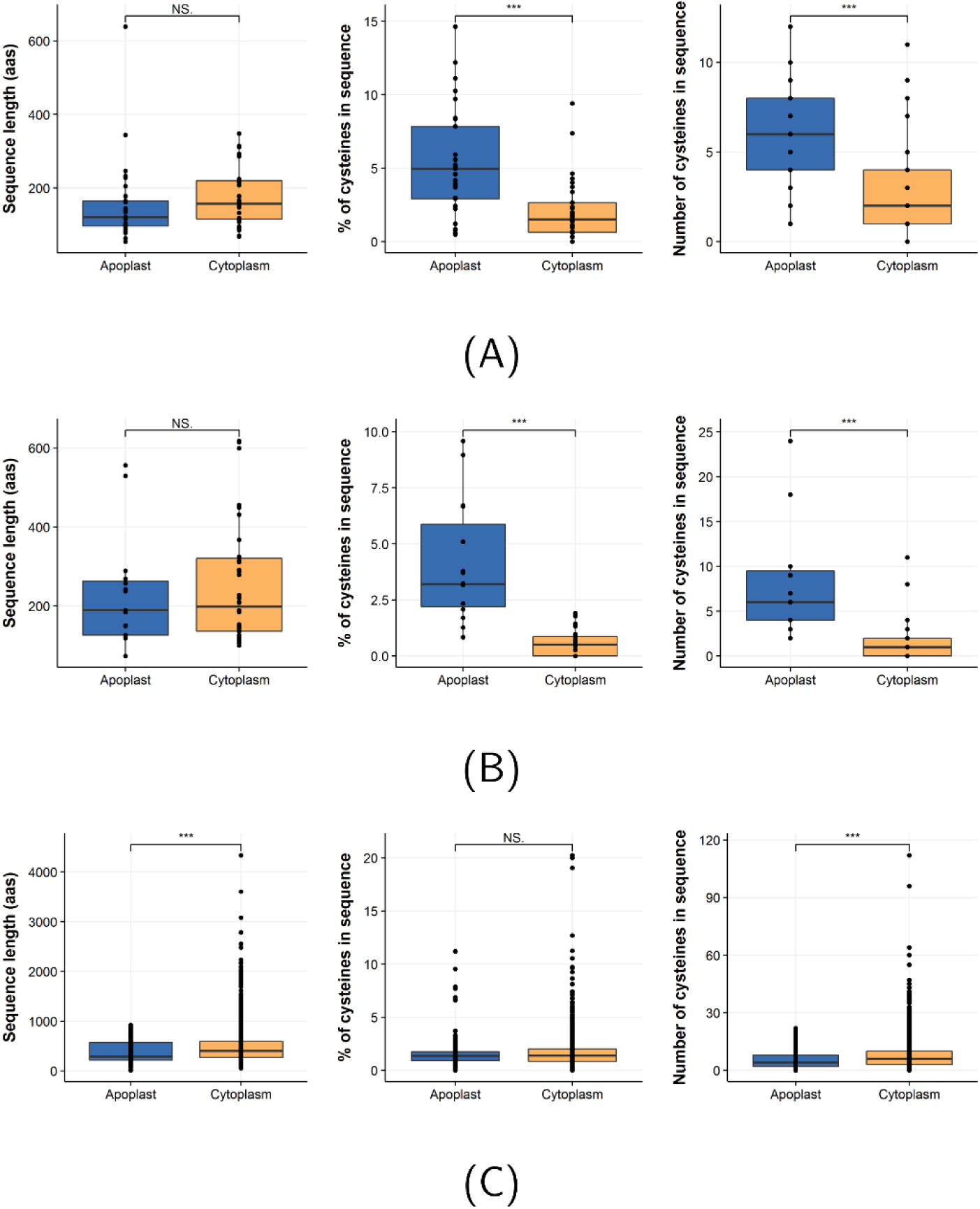
Differences in sequence length, percentage of cysteines and number of cysteines for apoplastic and intracellular (cytoplasmic) proteins. **(A)** Fungal apoplastic and cytoplasmic effectors. **(B)** Oomycete apoplastic and cytoplasmic effectors. **(C)** Apoplastic plant proteins and cytoplasmic plant proteins. All data points were drawn on top of the box plots.

**Table 2:**
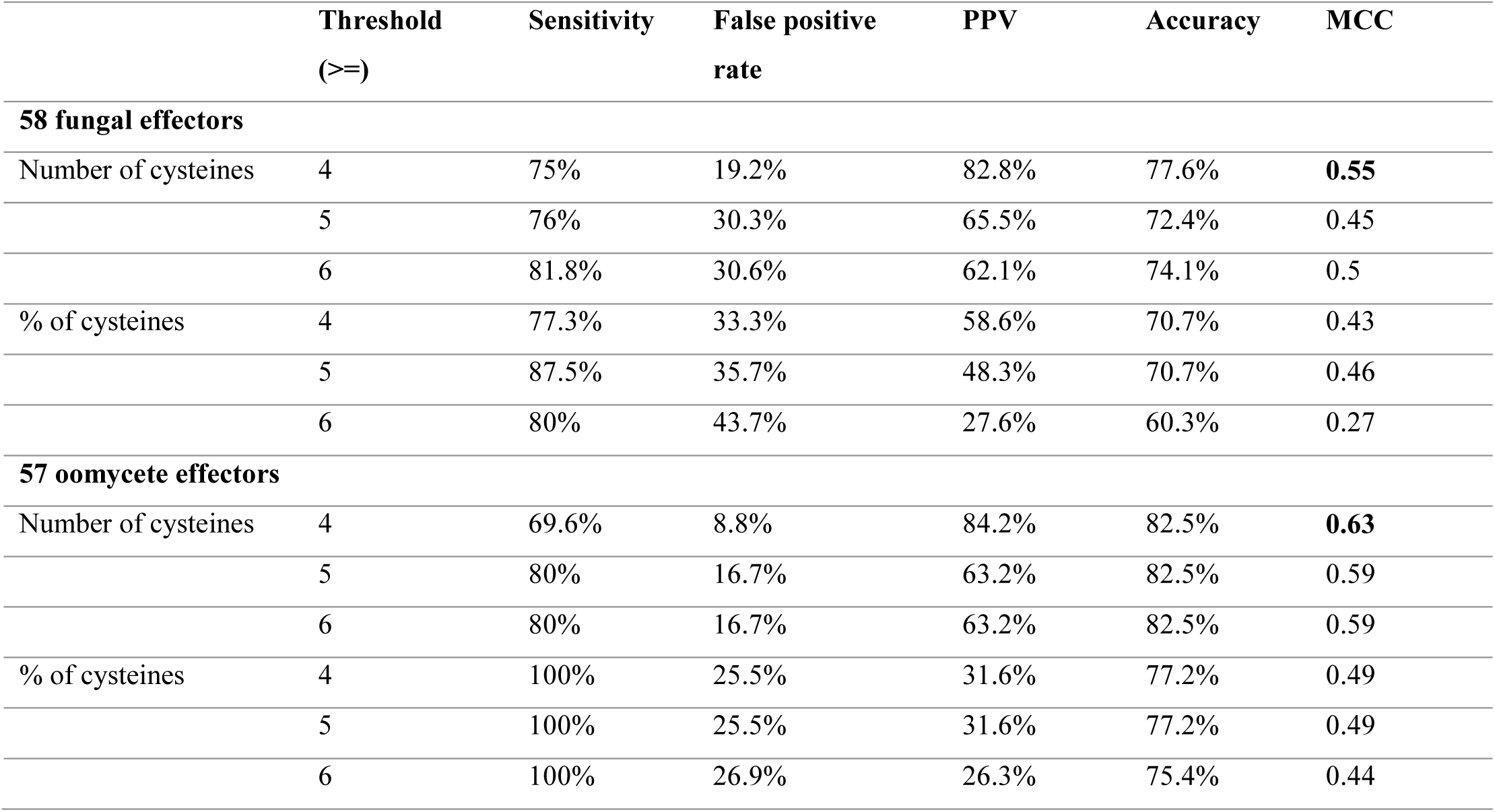
Performance of simple classifiers that predict apoplastic effectors based on cysteine residues. PPV stands for positive predictive value and MCC for Matthews Correlation Coefficient MCC.

|  | Threshold<br>(>=) | Sensitivity | False positive<br>rate | PPV | Accuracy | MCC |
| --- | --- | --- | --- | --- | --- | --- |
| **58 fungal effectors** |  |  |  |  |  |  |
| Number of cysteines | 4 | 75% | 19.2% | 82.8% | 77.6% | **0.55** |
|  | 5 | 76% | 30.3% | 65.5% | 72.4% | 0.45 |
|  | 6 | 81.8% | 30.6% | 62.1% | 74.1% | 0.5 |
| % of cysteines | 4 | 77.3% | 33.3% | 58.6% | 70.7% | 0.43 |
|  | 5 | 87.5% | 35.7% | 48.3% | 70.7% | 0.46 |
|  | 6 | 80% | 43.7% | 27.6% | 60.3% | 0.27 |
| **57 oomycete effectors** |  |  |  |  |  |  |
| Number of cysteines | 4 | 69.6% | 8.8% | 84.2% | 82.5% | **0.63** |
|  | 5 | 80% | 16.7% | 63.2% | 82.5% | 0.59 |
|  | 6 | 80% | 16.7% | 63.2% | 82.5% | 0.59 |
| % of cysteines | 4 | 100% | 25.5% | 31.6% | 77.2% | 0.49 |
|  | 5 | 100% | 25.5% | 31.6% | 77.2% | 0.49 |
|  | 6 | 100% | 26.9% | 26.3% | 75.4% | 0.44 |

For 19 experimentally validated apoplastic oomycete effectors and 38 experimentally validated cytoplasmic oomycete effectors (Table 1), we observed no significant differences in sequence length distribution (Fig. 1B). However, apoplastic oomycete effectors are significantly enriched in cysteines compared to cytoplasmic oomycete effectors. We tested different thresholds for cysteine content and found that a threshold of >= four cysteines achieved sensitivity of 69.6% and false positive rate of 8.8%. This suggests that a simple classifier using a threshold of at least four cysteines in the sequence can predict oomycete apoplastic effectors more accurately than fungal apoplastic effectors (Table 2). However, there are exceptions such as the oomycete pathogen *Phytophthora sojae* that employs an essential apoplastic effector called PsXEG1 with only two cysteines in its sequence (Ma *et al.*, 2015).

We then compared the distribution of sequence length and cysteine content for 349 apoplastic plant proteins and 1,950 intracellular plant proteins (see Methods). Apoplastic plant proteins have significantly shorter sequences and a lower number of cysteines compared to intracellular plant proteins (Fig. 1C). For example, the intracellular plant proteins MT2C, a metallothioneine-like potein from *Oryza sativa subsp. japonica* (UniProt entry A3AZ88) is localized in the cytosol yet has a sequence length of 84 aas and 17 cysteines and the intracellular transcriptional regulator NFXL2 from *Arabidopsis thaliana* (UniProt entry Q9FFK8) has 112 cysteines and sequence length of 883 aas. Taken together, we conclude that neither a small size nor high cysteine content alone are discriminative features for predicting apoplastic localization of both plant and effector proteins. In the following, we use machine learning to investigate if additional protein properties determine if a protein localizes to the plant apoplast.

### Training of a machine learning classifier for predicting effector and plant protein localization to the apoplast

To assess if protein properties can accurately distinguish apoplastic proteins from cytoplasmic proteins for both effectors and plant proteins, we trained a machine learning classifier (Fig. 2). We combined apoplastic plant proteins and randomly selected fungal and oomycete effector proteins (Table 1) as positive training data and intracellular plant proteins as negative training data. Both positive and negative training data were homology-reduced for training the machine learning classifier (see Methods). We deliberately did not remove the signal peptides for the secreted, apoplastic proteins in the positive training set because the state-of-the-art signal peptide cleavage site prediction software SignalP (Petersen *et al.*, 2011) does not allow reuse or incorporation of the code into other programs. An alternative to incorporating SignalP automatically into ApoplastP is a manual removal of e.g. the first 20 aas as the default signal peptide region. However, this would also remove N-termini of non-secreted proteins and of Golgi-independent secreted proteins lacking a signal peptide. Requiring users to submit sequences where signal peptides have been taken off requires scripting skills for parsing the outputs of SignalP, especially for versions prior to SignalP 4.1 that are the most sensitive for effector signal peptide prediction (Sperschneider *et al.*, 2015b). We later show that the inclusion of the signal peptide region in the positive training data has minimal effect on the performance of the machine learning classifier.

**Fig. 2:**
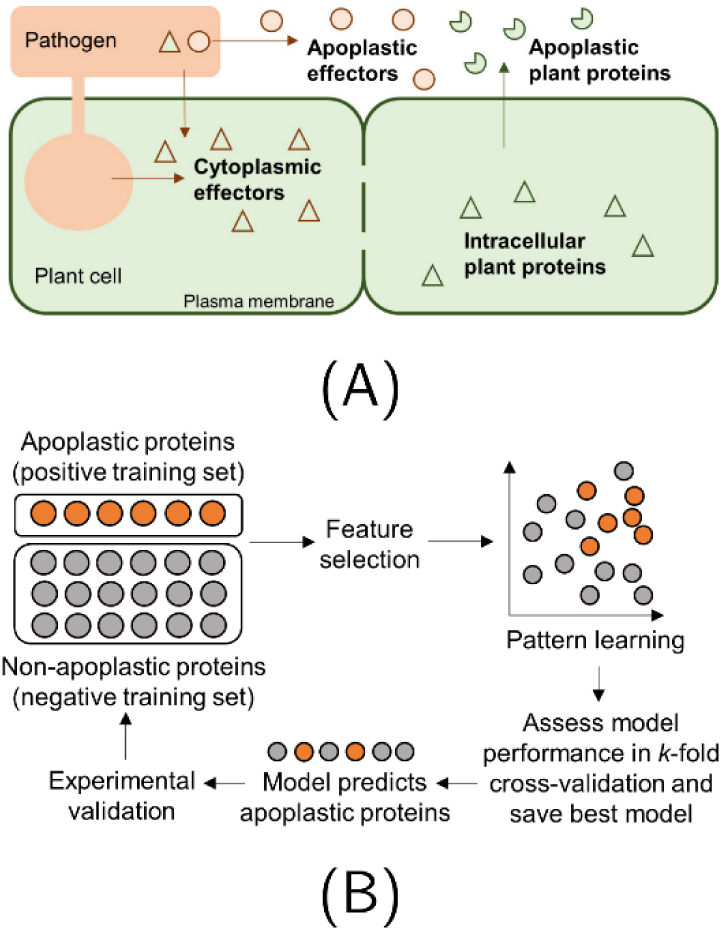
Training of a machine learning classifier for apoplastic protein prediction. **(A)** Apoplastic effectors and apoplastic plant proteins were used as positive training data and intracellular plant proteins as negative training data. Cytoplasmic effectors were used as an independent test set. **(B)** Positive and negative training data are used to train machine learning classifiers using selected features. A common technique of assessing performance is to use *k*-fold cross-validation, which can assess how a classifier is able to generalize to an independent dataset. In *k*-fold cross-validation, the training data are partitioned into *k* sets of equal size. The classifier is trained on *k*–1 datasets and tested on the one holdout set. This procedure is repeated *k* times and performance is reported. The best model is saved for effector prediction. Predicted apoplastic proteins can be taken to experimental validation and can be included in re-training of the classifier for improved classification in the future.

The homology-reduced negative training data are significantly larger than the homology-reduced positive training data (1,773 proteins compared to 84 proteins) and therefore, randomly selected smaller sets from the negative training data were chosen in the training of the classifier (100 sets generated for each of the ratios between the number of positive and negative training examples of 1:3, 1:4, 1:5, resulting in 300 negative sets varying in size from 252 to 420 proteins). For each protein, a feature vector was calculated using amino acid frequencies, amino acid class frequencies, number of cysteines, protein net charge, isoelectric point, grand average of hydrophobicity (GRAVY), protein instability index and aromaticity. We assessed the average performance of various machine learning classifiers and found that overall, the Random Forest classifier performed best (data not shown). We chose the best Random Forest model ranked in terms of AUC (area under the ROC curve) amongst all 300 trained models as the classifier called ApoplastP (ratio 1:4). In 10-fold cross-validation, ApoplastP achieves sensitivity of 58.3%, specificity of 98.2%, PPV of 89.1%, MCC of 0.67 and AUC of 0.951. As the 10-fold cross-validation is predominantly evaluated on plant proteins, we also directly compared the performance of ApoplastP to the simple classifiers based on cysteine thresholds from the previous section. On the set of effectors that share no overlap with the training data, ApoplastP improves accuracy by 12.5% for fungi and by 13.7% for oomycetes (Table 3).

**Table 3:** Performance of simple classifiers that predict apoplastic effectors based on cysteine residues compare to ApoplastP. PPV stands for positive predictive value and MCC for Matthews Correlation Coefficient MCC.

| Classifier | Sensitivity | False positive<br>rate | PPV | Accuracy | MCC |
| --- | --- | --- | --- | --- | --- |
| **58 fungal effectors** |  |  |  |  |  |
| Cysteines >=4 | 75% | 19.2% | 82.8% | 77.6% | 0.55 |
| ApoplastP | **89.7%** | **10.3%** | **86.7%** | **89.7%** | **0.79** |
| **57 oomycete effectors** |  |  |  |  |  |
| Cysteines >=4 | 69.6% | 8.8% | 84.2% | 82.5% | 0.63 |
| ApoplastP | **89.5%** | **0%** | **100%** | **96.5** | **0.92** |
| **40 fungal effectors (no overlap with ApoplastP training set)** |  |  |  |  |  |
| Cysteines >=4 | 50% | 12.5% | **72.7%** | 72.5% | 0.41 |
| ApoplastP | **72.7%** | **10.3%** | **72.7%** | **85%** | **0.62** |

| 51 oomycete effectors (no overlap with ApoplastP training set) |  |  |  |  |  |
| --- | --- | --- | --- | --- | --- |
| Cysteines >=4 | 61.6% | 6.1% | 84.6% | 82.4% | 0.6 |
| ApoplastP | **84.6%** | **0%** | **100%** | **96.1%** | **0.9** |

We selected the six most discriminative features that separate non-apoplastic from apoplastic proteins as predicted by WEKA and plotted their distribution in the positive and training sequence data. Overall, apoplastic proteins appear to be enriched in small amino acids, tiny amino acids and cysteines as well as depleted in glutamic acid, charged amino acids and acidic amino acids (Fig. 3). The enrichment and depletion analysis confirms that apoplastic localization is not a feature of a high cysteine content alone and that machine learning is sensitive to discovering compositional patterns of apoplastic proteins.

**Fig. 3:**
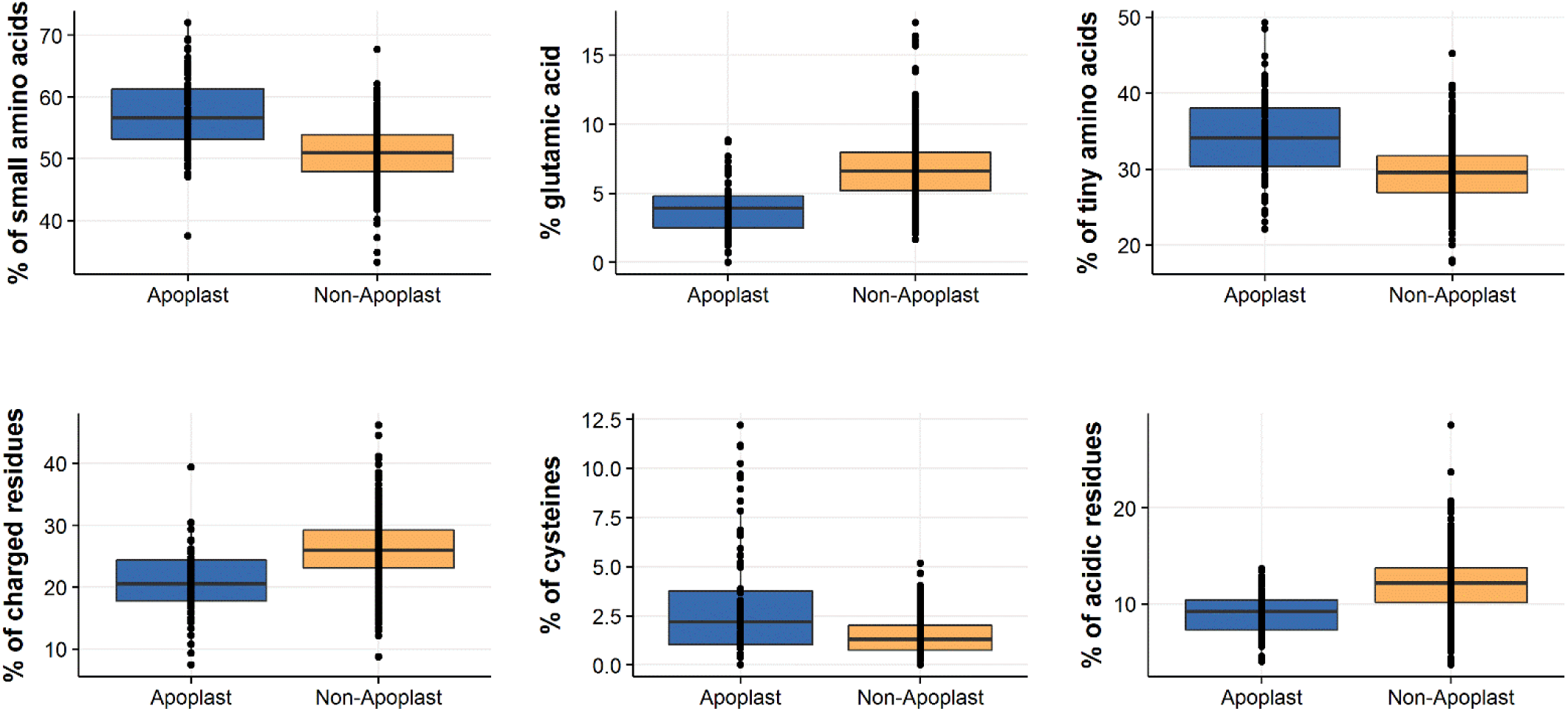
Box plots of feature distribution for the ApoplastP training set. The positive training set consists of 84 apoplastic plant and effector proteins and the negative training set consists of 336 non-apoplastic plant proteins. All data points were drawn on top of the box plots.

### The signal that separates apoplastic proteins from non-apoplastic proteins is not related to the presence of a signal peptide

As the positive training set consists of protein sequences with signal peptides and the negative training set consists of protein sequences without a signal peptide, we first assessed if ApoplastP is biased towards recognizing properties relating to secretion alone. Thus, we tested ApoplastP on secreted proteins (including their signal peptides) that do not reside in the plant apoplast. The first set we used is cytoplasmic effectors as these are secreted but enter the plant cell and act intracellularly (Table 1). ApoplastP correctly predicts all 38 experimentally validated cytoplasmic oomycete effectors (RXLR effectors and Crinklers) as nonapoplastic. For the 29 experimentally validated cytoplasmic fungal effectors, ApoplastP returns three false positives (10.3% false positive rate), all from *Magnaporthe oryzae* (AvrPiz-t, Avr-Pii, Avr-Pik). AvrPiz-t and Avr-Pik are part of the MAX (***M****agnaporthe* **A**vrs and To**x**B like) effector family of sequence-unrelated but structurally conserved fungal effectors (de Guillen *et al.*, 2015). The MAX effector family member ToxB is an effector that is secreted into the apoplast and acts extracellularly (Figueroa *et al.*, 2015) and the similarity on the structure level to the intracellular effector AvrPiz-t and Avr-Pik could explain their prediction as apoplastic. Taken together, we estimate that ApoplastP has a false positive rate of 4.4% on cytoplasmic effectors, as compared to 1.8% in 10-fold cross-validation on intracellular plant proteins. The removal of the first 20 aas as the default signal peptide region has no impact on the false positive rate for this set (Table 4). ApoplastP also has a low false positive rate of 0.8% on 358 RXLR effector candidates (HMM model, Win *et al.* (2007)).

**Table 4:**
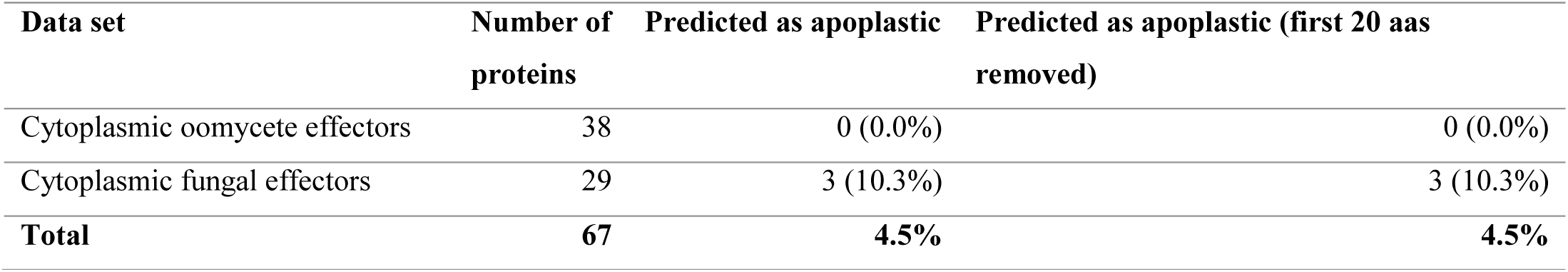
Independent test set consisting of secreted, cytoplasmic effectors.

| Data set | Number of proteins | Predicted as apoplastic | Predicted as apoplastic (first 20 aas removed) |
| --- | --- | --- | --- |
| Cytoplasmic oomycete effectors | 38 | 0 (0.0%) | 0 (0.0%) |
| Cytoplasmic fungal effectors | 29 | 3 (10.3%) | 3 (10.3%) |
| **Total** | **67** | **4.5%** | **4.5%** |

We then used non-apoplastic fungal, plant and mammalian proteins with a predicted signal peptide to further assess the false positive rate of ApoplastP. Proteins with a predicted signal peptide are not necessarily released to the extracellular space, but can be retained in the endoplasmic reticulum (ER) or Golgi apparatus, be directed to the lysosome or vacuole, contain transmembrane helices or a GPI-anchor that anchors it to the outer face of the plasma membrane. We took plant and fungal proteins that have been experimentally shown to localize to the ER, Golgi, vacuole or contain transmembrane domains, yet also have a predicted signal peptide. Plant GPI-anchored proteins can be anchored to the apoplastic face of the membranes and those from pathogens have been found to interact with host cells and can be required for virulence, therefore we did not include them. We also took extracellular mammalian proteins with a predicted signal peptide as a negative test set. Overall, ApoplastP has a false positive rate of 6% on all 1,217 plant, fungal and mammalian non-apoplastic proteins with a predicted signal peptide (Table 5). We observed the highest false positive rate (16.1%) on the set of plant proteins localized to the vacuole. The five mis-predicted plant proteins are annotated; three endochitinases, a hevein-like preproprotein with putative antimicrobial activities and a glycine-rich protein. The three endochitinases in particular are annotated in UniProt as involved in defense against chitin-containing fungal pathogens, which indicates that the localization to the vacuole is either a mis-annotation or that they are indeed apoplastic proteins that are directed to the vacuole for storage and released upon infection. The set of ER-localized fungal proteins also has a high false positive rate of 14.1% and 8 out of 10 mis-predicted proteins are annotated as uncharacterized proteins from *Schizosaccharomyces pombe*. The removal of the first 20 aas as the default signal peptide region increases the false positive rate to 8.0% on the overall set.

**Table 5:**
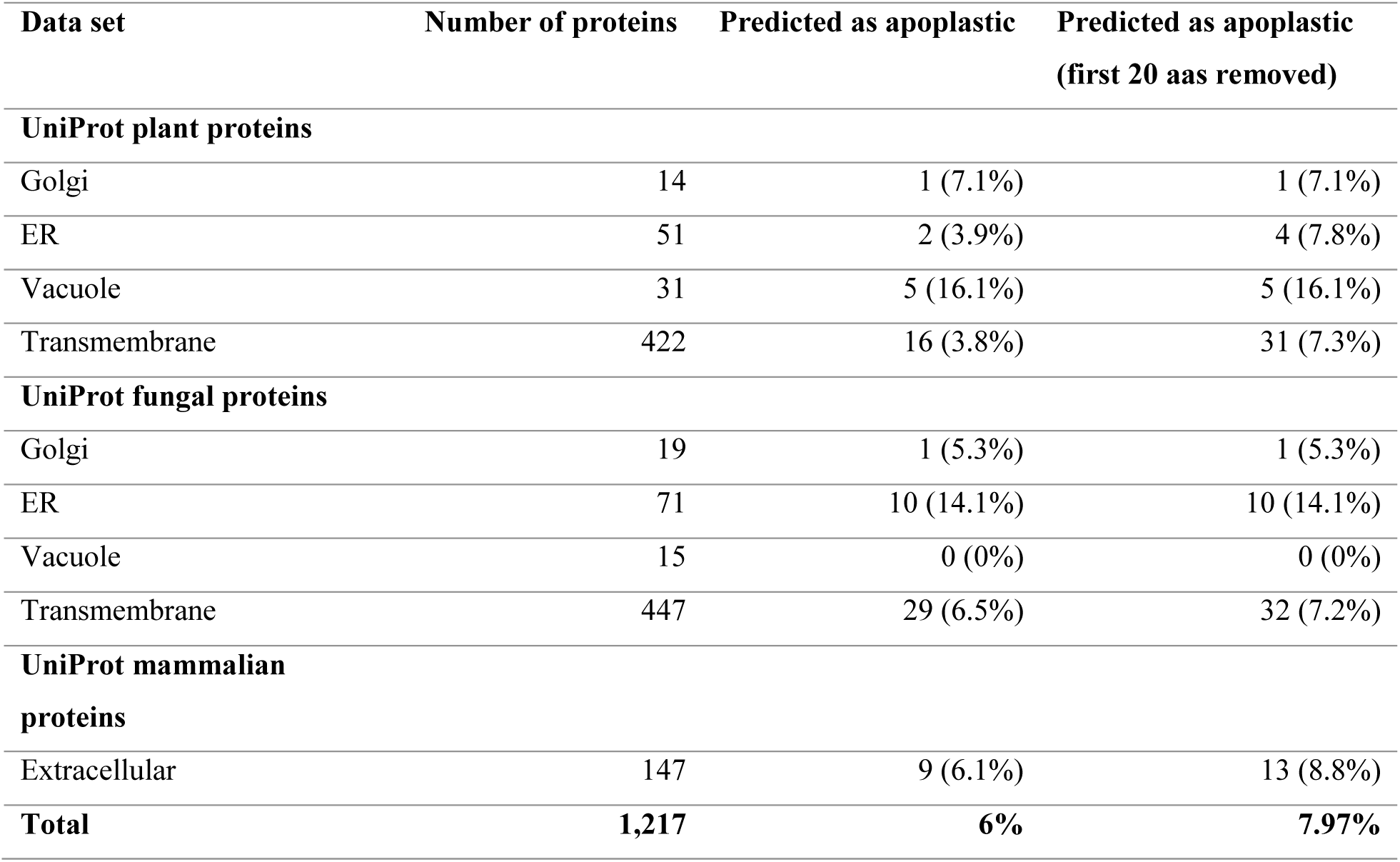
Independent test set consisting of non-apoplastic proteins with a predicted signal peptide.

Secreted proteins conventionally carry a signal peptide and enter the ER/Golgi pathway before being released to the extracellular space. Unconventional secretion of proteins lacking a signal peptide has also been reported and is commonly induced by stress (Rabouille, 2017). However, experimental identification of leaderless secretion is technically challenging and currently, there is only one example in plants of a protein with a positive immunolocalization in the apoplast, namely a lectin from sunflower (*Helianthus annuus*) (Pinedo *et al.*, 2012). For this particular study by Pinedo *et al.* (2012), ApoplastP predicts only 1 out of 14 proteins identified from extracellular fluid as apoplastic, namely the apoplast-localized lectin. (Table 6). The other 13 proteins are annotated as a golgi-membrane localized hexosyltransferase, a cytochrome p450 protein, a mitochondrial pentatricopeptide protein, a splicing factor Sc35 protein, an amidase protein, a mitochondrial maturase protein, a mutator like-transposase, an LEA protein, a membrane-localized heat shock protein, a transcription factor, an embryonic DC-8 protein, a WEB family protein and a protein kinase protein, which indicates their likely localization to membranes or the plant intracellular space.

**Table 6:**
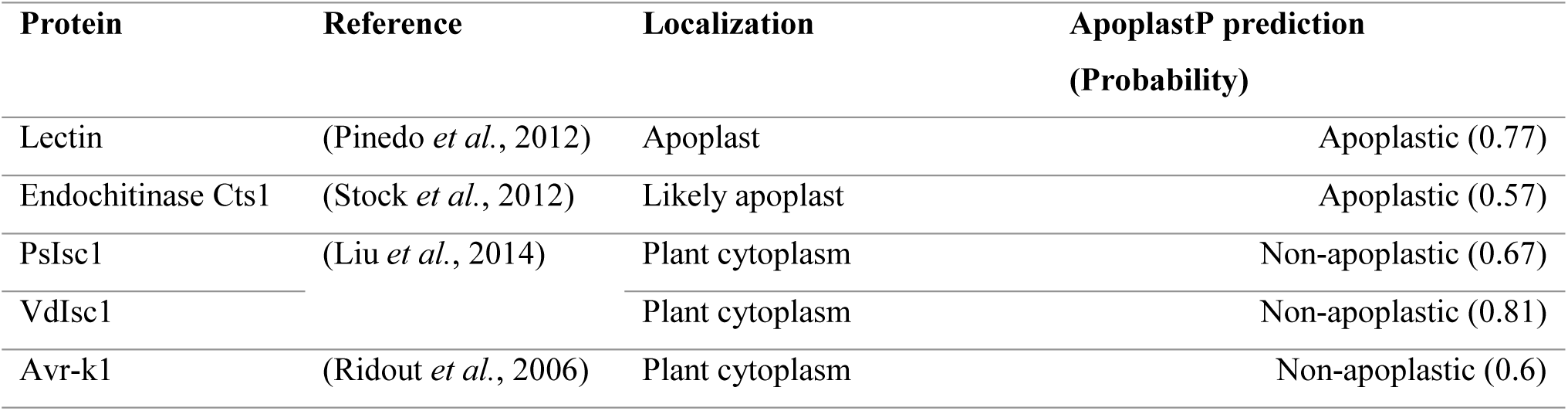
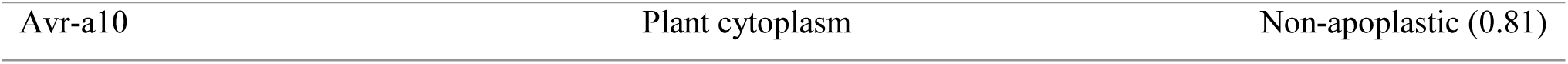
Unconventionally secreted proteins from plants and fungi with experimental validation.

| Protein | Reference | Localization | ApoplastP prediction (Probability) |
| --- | --- | --- | --- |
| Lectin | (Pinedo *et al.*, 2012) | Apoplast | Apoplastic (0.77) |
| Endochitinase Cts1 | (Stock *et al.*, 2012) | Likely apoplast | Apoplastic (0.57) |
| PsIsc1 | (Liu *et al.*, 2014) | Plant cytoplasm | Non-apoplastic (0.67) |
| VdIsc1 |  | Plant cytoplasm | Non-apoplastic (0.81) |
| Avr-k1 | (Ridout *et al.*, 2006) | Plant cytoplasm | Non-apoplastic (0.6) |

|  |  |  |
| --- | --- | --- |
| Avr-a10 | Plant cytoplasm | Non-apoplastic (0.81) |

Unconventionally secreted proteins from fungi include the Cts1 endochitinase from *Ustilago maydis* with a putative apoplastic localization and this is also predicted as apoplastic by ApoplastP. The isochorismatase effectors PsIsc1 and VdIsc1 from *Phytophthora sojae* and *Verticillium dahliae*, respectively, (Liu *et al.*, 2014) as well as the *Blumeria graminis* f. sp. *hordei* effectors Avr-k1 and Avr-a10 are thought to be unconventionally secreted, although localized to the plant cytoplasm. ApoplastP correctly predicts these four secreted cytoplasmic effectors as non-apoplastic. Finally, we applied ApoplastP to the full *Arabidopsis thaliana* proteome and 1,938 of 27,426 proteins (7.1%) are predicted as apoplastic. SignalP 4.1 predicts a signal peptide for 60.2% of the 1,938 putative apoplastic proteins.

Lastly, we used RNA sequencing data for germinated spores (urediniospores) and haustorial tissue from the wheat stem rust fungus *Puccinia graminis* f. sp. *tritici* 21-0 (Upadhyaya *et al.*, 2014) and performed differential expression analysis. For genes with high up-regulation in haustoria that encode secreted proteins, ApoplastP predicts only 9.1% as apoplastic (Table 7). This is consistent with the haustorial structure in rust fungi, in which the extra-haustorial matrix is thought to be separated from the plant apoplast by a neckband and the role of haustoria as the main site of cytoplasmic effector delivery (Voegele & Mendgen, 2003; Garnica *et al.*, 2014). In contrast, a simple classifier using a threshold of at least four cysteines as a criterion for apoplastic localization returns 30.9% of secreted proteins that are encoded by genes with high up-regulation in haustoria as apoplastic. This confirms the high false positive rate of cysteine-rich classifiers for apoplastic localization prediction observed in the previous sections.

**Table 7:** ApoplastP prediction results on secreted proteins from *P. graminis f. sp. tritici*.

| Condition | log FC | Number of genes | ApoplastP predicted as apoplastic | Cysteine-rich classifier ($\geq 4$) |
| --- | --- | --- | --- | --- |
| Up-regulated in haustoria vs. germinated spores | $\geq 1.0$ | 791 | 158 (20.0%) | 52.7% |
| | $\geq 10$ | 55 | 5 (9.1%) | 30.9% |
| Up-regulated in germinated spores vs. haustoria | $\geq 1.0$ | 458 | 145 (31.7%) | 55.9% |
| | $\geq 10$ | 26 | 12 (46.2%) | 65.4% |

### ApoplastP correctly identifies 75% of apoplastic effectors in independent test sets

We used an independent test set of 32 apoplastic effectors from fungi, oomycetes and nematodes to assess the true positive rate (correctly identified apoplastic proteins) of ApoplastP. We found that ApoplastP delivers a high true positive rate of 75% on the experimentally validated apoplastic effectors, but does not identify 8 effectors (AvrLm1, PstSCR1, CfTom1, EPI10, OPEL, Crt-1, HYP-3 and CLE-1) as apoplastic (Table 8).

**Table 8:**
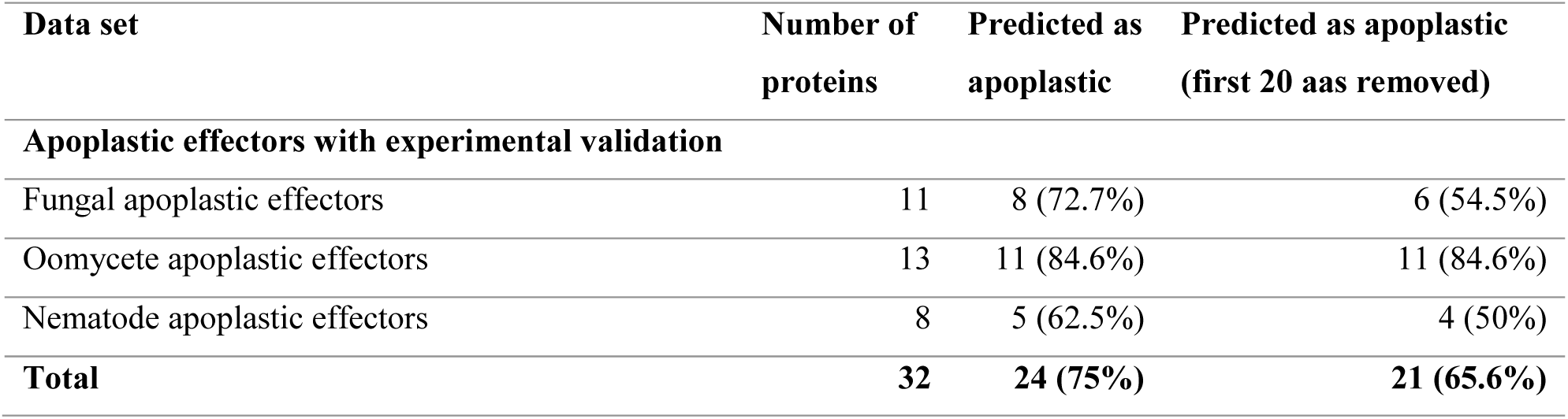
Independent apoplastic effector test sets.

We then tested ApoplastP on 923 apoplastic proteins from both plant and pathogens that were determined using proteomics (Table 9). Apoplastic proteomics is prone to false positives due to the potential for cell damage that can lead to contamination of the sample with cytoplasmic proteins (Delaunois *et al.*, 2014). Therefore, we tested ApoplastP using both the apoplastic proteome set as well as on only the 480 proteins in these sets that have a predicted signal peptide using SignalP 4.1 (Petersen *et al.*, 2011). We observed the lowest number of predicted apoplastic proteins (23.8%) in the *Magnaporthe oryzae* apoplastic proteome during rice infection (Kim *et al.*, 2013) and the highest number of predicted apoplastic proteins (80%) in the apoplastic proteome of *Nicotiana benthamiana* leaves (Goulet *et al.*, 2010), with an average prediction rate on all proteomics sets of 33%. Applying ApoplastP to only the proteins with a predicted signal peptide increases the prediction rate to an average of 55.2%. In the previous section we showed that ApoplastP correctly predicts the localization of six unconventionally secreted proteins and despite this being a small test set, it could indicate that the proteomics sets do indeed contain substantial contamination from cytoplasmic proteins or cell wall proteins.

**Table 9:** Apoplastic proteomics test sets.

| Data set | SignalP 4.1 predicted as secreted | Number of proteins | Predicted as apoplastic |
| --- | --- | --- | --- |
| Extracellular proteome of *P. infestans* (Meijer *et al.*, 2014) | - | 199 | 100 (50.3%) |
|  | Yes | 180 | 95 (52.8%) |
| *Magnaporthe* apoplastic proteome (Kim *et al.*, 2013) | - | 403 | 96 (23.8%) |
|  | Yes | 155 | 81 (52.3%) |
| Rice apoplastic proteome (Kim *et al.*, 2013) | - | 249 | 68 (27.3%) |
|  | Yes | 94 | 53 (56.4%) |
| Leaf apoplast proteome of *Brassica napus* var. *napus* after infection with *Verticillium longisporum* (Floerl *et al.*, 2008) | - | 9 | 4 (44.4%) |
|  | Yes | 8 | 4 (50%) |
| Leaf apoplast proteome of *Arabidopsis thaliana* after infection with *Verticillium longisporum* (Floerl *et al.*, 2012) | - | 43 | 21 (48.8%) |
|  | Yes | 27 | 19 (70.4%) |
| Apoplastic proteome of *Nicotiana benthamiana* leaves (Goulet *et al.*, 2010) | - | 20 | 16 (80%) |
|  | Yes | 16 | 13 (81.3%) |
| **Total** | **-** | **923** | **305 (33%)** |
|  | **Yes** | **480** | **265 (55.2%)** |

### The secretomes of saprophytic fungi, necrotrophic plant pathogens and extracellular fungal pathogens are enriched for predicted apoplastic proteins

We applied ApoplastP to the predicted secretomes of published fungal and oomycete genomes (see Methods) and plotted the percentages of predicted apoplastic proteins (Fig. 4A). Overall, the proportions of predicted apoplastic proteins in secretomes correspond well with the extracellular and intracellular colonization strategies of the fungal and oomycete pathogens that were tested. The highest proportions of predicted apoplastic proteins were recorded for the secretomes of the wood rotting saprophyte *Dichomitus squalens* (57.3%), the white rot saprophytes *Punctularia strigosozonata* (57.3%) and *Stereum hirsutum* (57.2%), followed by the broad host range necrotrophic fungal pathogens *Sclerotinia sclerotiorum* (55.8%) and *Botrytis cinerea* (55.7%). The lowest proportions of predicted apoplastic proteins were recorded for the secretomes of the obligate biotrophic oomycete pathogens *Albugo laibachii* (10%) and *Hyaloperonospora arabidopsidis* (15.2%), the plant-pathogenic yeast *Ashbya gossypii* (18.1%), the animal pathogen *Batrachochytrium dendrobatidis* (20%) and the obligate biotrophic fungal pathogens *Blumeria graminis* f. sp. *tritici* (22.9%) and *B. graminis* f. sp. *hordei* (23.1%). In the following, we labelled *Venturia pirina, V. inaequalis, C. fulvum, L. maculans* and *Z. tritici* as apoplastic fungal pathogens and removed *L. maculans* and *Z. tritici* from the set of hemibiotrophic fungal pathogens. We then compared groups containing at least two species to fungal saprophytes. Compared to fungal saprophytes, the percentage of predicted apoplastic proteins in the secretome is significantly lower for obligate biotrophic pathogens, hemibiotrophic pathogens and fungal plant symbionts, but not for apoplastic fungal pathogens or necrotrophic fungal pathogens (Fig. 4B).

**Fig. 4:**
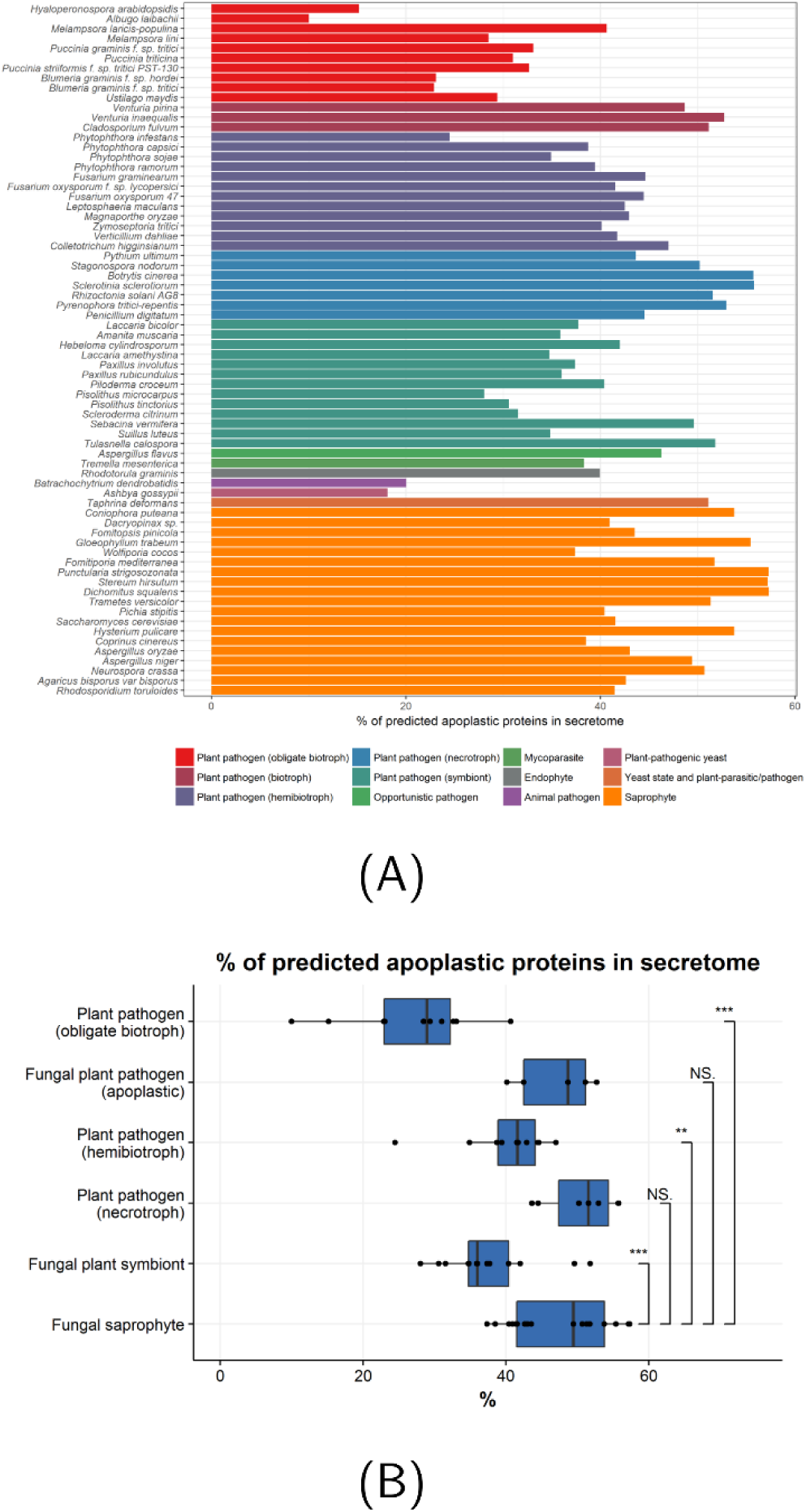
**(A)** Percentages of ApoplastP predicted apoplastic proteins in secretomes of fungi and oomycetes. **(B)** Box plots of predicted apoplastic proteins in secretomes grouped according to lifestyle. All data points were drawn on top of the box plots.

### Apoplastic proteins are highly enriched for predicted fungal effectors in rust pathogens

Next, we assessed the proportion of predicted apoplastic and non-apoplastic effector proteins using EffectorP (Sperschneider *et al.*, 2016). We did not apply EffectorP to the oomycete secretomes as it is designed specifically for fungal secretomes. The highest percentages of predicted effectors in the apoplastic protein set was recorded for the rust pathogens *Puccinia striiformis* f. sp. *tritici* PST-130 (61.2%), *Puccinia graminis* f. sp. *tritici* (61.6%) and *Melampsora laricis-populina* (58.7%), whereas the lowest percentages were recorded for the fungal saprophytes *Pichia stipitis* (8.4%), *Hysterium pulicare* (8.9%) and *Wolfiporia cocos* (9%). We compared the percentages of predicted effectors in the apoplastic set to predicted effectors in the non-apoplastic set (Fig. 5). Amongst pathogenic fungi, we found significant differences only for the rust pathogens, with an average of 52.1% apoplastic proteins predicted as effectors, whereas only 33.5% of non-apoplastic secreted proteins are predicted as effectors. An outlier in this set is *Melampsora lini*, which has only 25.7% apoplastic proteins predicted as effectors. Taken together, this indicates that the prediction abilities of EffectorP and ApoplastP are distinct, and that whilst the percentage of apoplastic proteins in rust pathogen secretomes is low, they are highly enriched for predicted effectors.

**Fig. 5:**
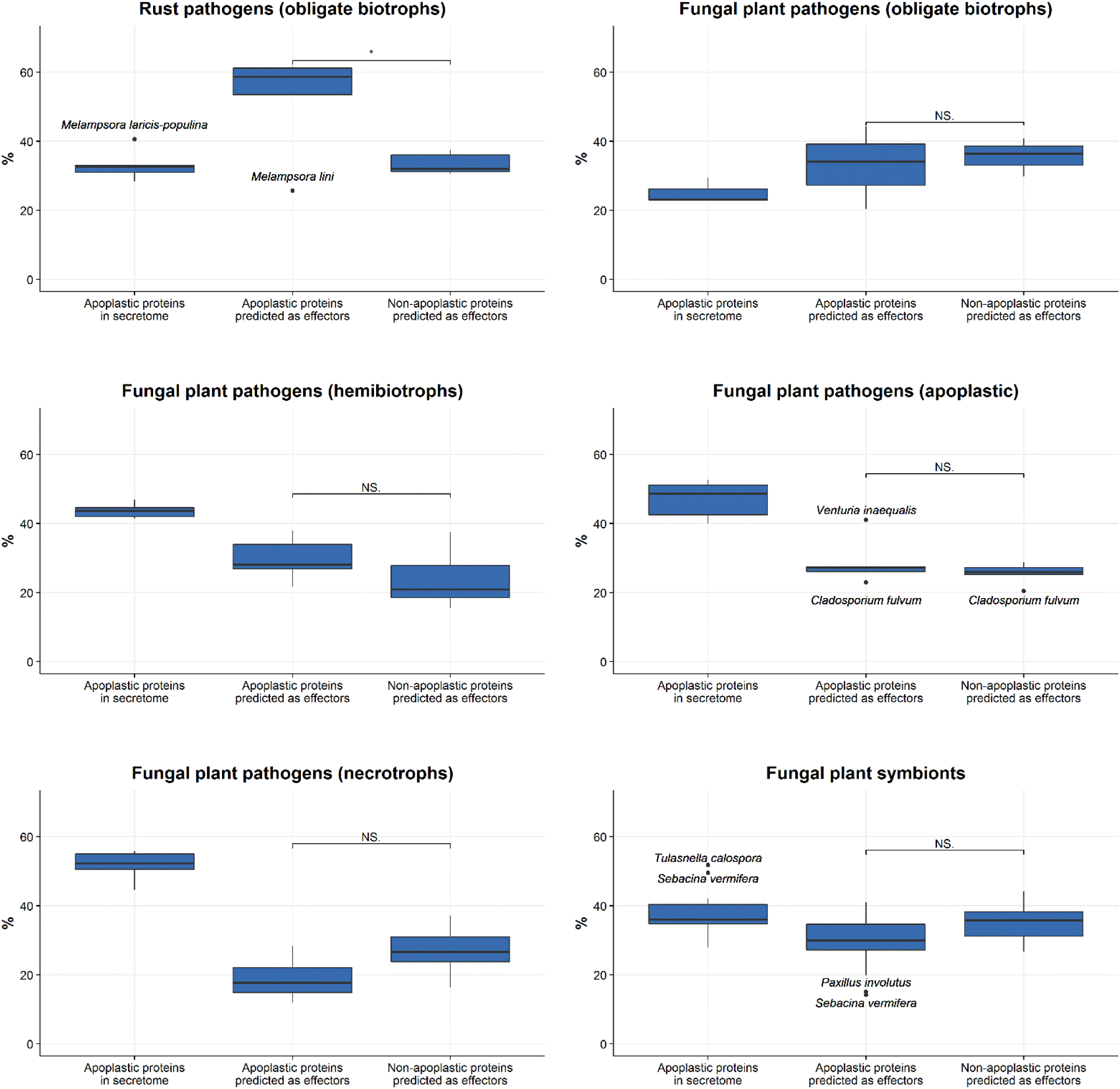
Percentages of EffectorP predicted effectors in the predicted apoplastic and non-apoplastic sets for fungi. Outliers were drawn and labelled around the box plots.

### Conserved sequence motifs in predicted cytoplasmic effector candidates

We predicted apoplastic and non-apoplastic (cytoplasmic) effector candidates in fungi using ApoplastP and EffectorP. To find conserved motifs in predicted cytoplasmic effector candidates, we reduced the sequence homology in each set and applied a MEME motif search (Bailey *et al.*, 2009) with the setting of one occurrence of a motif per sequence. Even though EffectorP is not designed for effector prediction in oomycetes, we used this methodology as a positive control on *Phytophthora infestans*. As expected, MEME returned the RxLR (yet with a non-significant E-value > 0.05) and dEER motifs (E-value 2.2x10^-28^) in the cytoplasmic effector candidate set (Fig. 6), but not in the apoplastic effector candidate set. For the fungal pathogens, we found the [YFW]xC motif in the predicted cytoplasmic effector candidate set of *Blumeria graminis* f. sp. *hordei* (E-value 1.2 x10^-33^, Fig. 6C), however it was also detected in the respective predicted apoplastic effector candidate set albeit with non-significant E-value (Fig. 6D). Weak conservation for the [YFW]xC motif was also found for the *Puccinia graminis* f. sp. *tritici* cytoplasmic effector candidate set (non-significant E-value > 0.05, Fig. 6E).

**Fig. 6:**
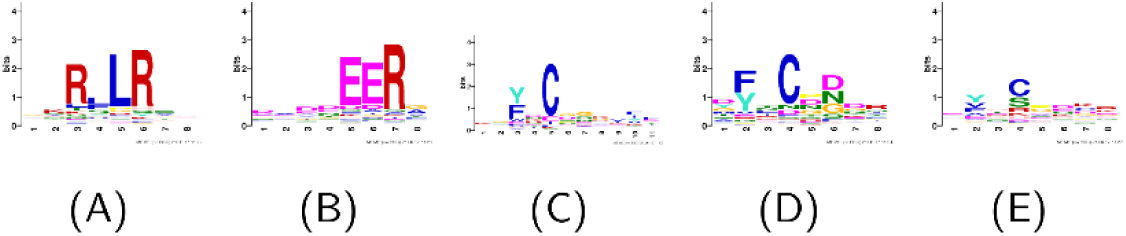
MEME motif searches on homology-reduced predicted apoplastic and non-apoplastic (cytoplasmic) effector proteins. For *Phytophthora infestans*, the RxLR (**A**) and dEER motifs (**B**) are predicted in the cytoplasmic effector candidates. In *Blumeria graminis* f. sp. *hordei*, the [YFW]xC motifs are predicted in both the non-apoplastic (**C**) and with weaker conservation in the cytoplasmic (**D**) effector candidate set. **(E)** The [YFW]xC motif is also found in the *Puccinia graminis* f. sp. *tritici* cytoplasmic effector candidate set with weaker conservation.

We also observed an enrichment in a proline at the +1 position after the predicted signal peptide cleavage site in fungal secretomes. We therefore performed a systematic search for +1 prolines in the mature protein sequences across the predicted secretomes of fungal and oomycete genomes using the predicted cleavage sites of the neural network model of SignalP 3 (Bendtsen *et al.*, 2004b). Whilst on average 9% of apoplastic plant proteins and 8.7% of oomycete secretomes have a +1 proline, this increases to 25.8% for fungal secretomes. For the fungal secretomes, 30.4% of predicted apoplastic proteins in fungal secretomes have a +1 proline compared to 21.3% of predicted non-apoplastic proteins. Significant differences between +1 proline content in predicted apoplastic and non-apoplastic proteins was observed for all fungal groups except obligate biotrophic fungal pathogens (Fig. 7). Furthermore, 34.5% of the 29 apoplastic fungal effectors and 41.4% of the 29 cytoplasmic fungal effectors have a +1 proline after the predicted signal peptide cleavage site. This includes the ToxA effector of *Pyrenophora tritici-repentis* and *Parastagonospora nodorum*, which has a pro-domain after the signal peptide region that is thought to be important for folding, but not necessary for toxic activity (Tuori *et al.*, 2000; Ciuffetti *et al.*, 2010). Taken together, this suggests that a +1 proline after the predicted signal peptide cleavage site is a prevalent characteristic of secreted fungal proteins, however it is unlikely related to fungal effector function.

**Fig. 7:**
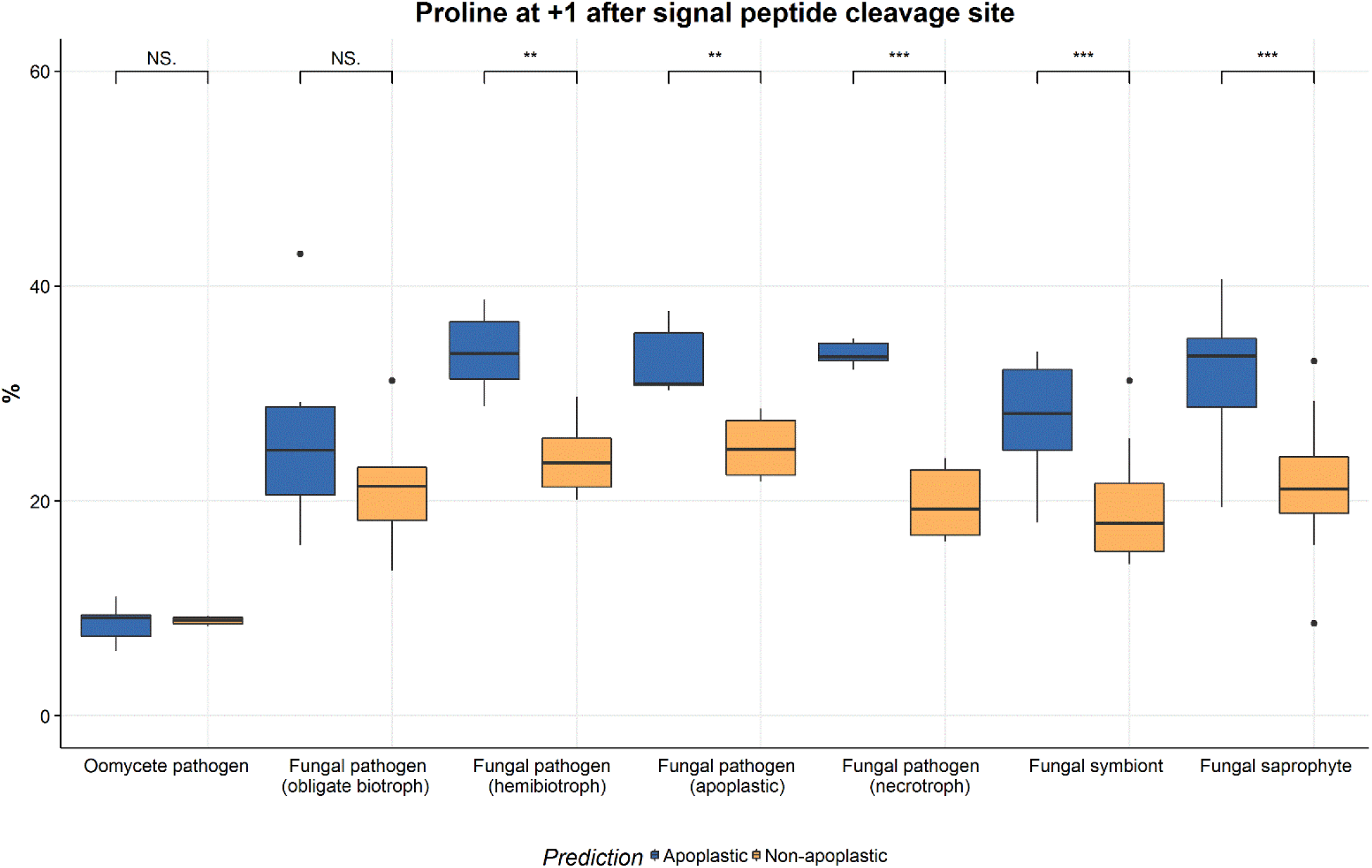
Percentages of proteins in fungal and oomycete secretomes that have a +1 proline after the signal 4 peptide cleavage site.

### Discussion

The plant apoplast is integral to essential plant processes such as intercellular signalling and transport. Furthermore, early interactions between plants and pathogens in the apoplast determine if a pathogen can colonize and infect plant tissue (Doehlemann & Hemetsberger, 2013). Whilst apoplastic localization prediction is important for effectors across all plant pathogen taxa as well as for secreted plant proteins, no dedicated computational method was previously available. Apoplastic proteins are commonly identified through microscopic analyses or apoplastic proteomics, however both techniques are technically challenging (Doehlemann & Hemetsberger, 2013; Delaunois *et al.*, 2014). Whilst tools such as SignalP (Petersen *et al.*, 2011) or Phobius (Kall *et al.*, 2004) can predict the presence of a signal peptide, proteins that are predicted to be secreted can also localize to the cell walls or be retained intracellularly (Emanuelsson *et al.*, 2007). Furthermore, effectors can either function in the plant apoplast or enter the plant cell cytoplasm and being able to discriminate between these two localizations accurately is highly desirable for shortlisting prime effector candidates for subsequent experimental validation.

Machine learning is a promising technique for effector prediction, because effectors co-localize with their respective plant targets and thus are likely to carry subcellular localization signals which may be cryptic. This also means that for training machine learning classifiers for effector localization prediction, one can take advantage of the large number of experimentally validated plant proteins with localization data, as effectors likely exploit the plant cell machinery for localization and function. Using both plant and effector localization data, we have pioneered a data-driven machine learning approach called ApoplastP that can predict if a protein localizes to the plant apoplast. By using machine learning, we were able to exploit compositional differences between apoplastic proteins and intracellular plant proteins that were previously unrecognized such as a depletion in glutamic acid for apoplastic proteins. ApoplastP outperforms the common approach of selecting apoplastic effectors from secretomes based on a high cysteine content, improving prediction accuracy by over 13%. For many pathogens, cytoplasmic effectors are delivered first to the plant apoplast and then subsequently enter plant cells, such as the SIX3, SIX5 and SIX6 effectors from *Fusarium oxysporum* f. sp. *lycopersici* (De Wit, 2016) or the ToxA effector from *Pyrenophora tritici-repentis* and *Parastagonospora nodorum* (Manning & Ciuffetti, 2005). We showed that ApoplastP recognizes the localization of cytoplasmic effectors with high accuracy, even if they enter the plant cell cytoplasm from the apoplast.

ApoplastP does not rely on the presence of a signal peptide and can thus predict unconventionally secreted proteins that localize to the plant apoplast. This makes it a valuable validation tool for screening apoplastic proteomics sets for cytoplasmic protein contamination and for elucidating unconventional secretion pathways in both plants and pathogens. Furthermore, ApoplastP will facilitate the identification of likely cytoplasmic effectors by exclusion and can potentially elucidate effector translocation mechanisms, e.g. through future compositional pattern searches in the predicted set of cytoplasmic effectors. In general, it highlights the benefit of using data-driven machine learning classifiers over classifiers that rely on user-driven thresholds in the field of plant-pathogen interactions.

## Acknowledgements

JS is supported by a CSIRO OCE Postdoctoral Fellowship. We thank Donald Gardiner and Jonathan Anderson for comments on earlier versions of this manuscript.

## Author contributions

J.S. planned and designed the research and developed the software. J.S, P.N.D., K.B.S. and J.M.T analysed data and wrote the manuscript.

